# Split GFP assay to show chloroplast targeting of *Agrobacterium* VirD2 protein

**DOI:** 10.1101/2023.12.14.571705

**Authors:** Aki Matsuoka, Julia L. Ferranti, Pal Maliga

## Abstract

Regenerating fertile *Arabidopsis thaliana* plants from tissue culture cells with transformed plastid genomes is difficult, because of somaclonal variation in tissue culture cells. For nuclear gene transformation, tissue culture limitations were overcome in Arabidopsis by direct transformation of the female gametes using the floral dip protocol and identification of transgenic events in the seed progeny. During *Agrobacterium* transformation the VirD2 protein guides the T-complex, consisting of single stranded transferred-DNA (T-DNA) coated with VirE2 proteins, to the plant nucleus. To enable floral dip transformation of the plastid DNA, we retargeted VirD2 to chloroplasts. We show plastid targeting of VirD2 in a split GFP assay, where VirD2-GFP_11_ complements GFP_1-10_ in chloroplasts. Floral dip transformation of plastids will avoid tissue culture altogether, making plastid transformation readily available for the research community.

## Introduction

High-frequency plastid transformation in Arabidopsis was achieved by choosing plants lacking a duplicated acetyl-CoA-carboxylase enzyme that makes plants hyper-sensitive to spectinomycin, the selective agent used in chloroplast transformation (Yu et al., 2017; Ruf et al., 2019; Yu et al., 2019). Inherent polyploidy of leaf cells and the tendency for somaclonal variation make difficult to regenerate fertile plants from transplastomic tissue culture cells (Galbraith et al., 1991; Melaragno et al., 1993; Bairu et al., 2011). Dipping Arabidopsis flowers in an *Agrobacterium* culture has made it possible to directly transform the female gametocyte without tissue culture and plant regeneration. Transgenic events have then been identified by germinating seeds on a selective medium (Clough and Bent, 1998; Desfeux et al., 2000; Zhang et al., 2006). Our goal is to enable *Agrobacterium*-mediated floral dip transformation of the plastid genome.

*Agrobacterium tumefacies* is a plant pathogen which transfers tumor-inducing genes to the plant nucleus (Lee and Gelvin, 2008; Lacroix and Citovsky, 2013). The tumor inducing plasmid has been disarmed by removing the tumor-inducing genes and the flanking T-DNA borders. The virulence (*vir*) genes were retained on a helper plasmid that encodes all functions required for T-DNA transfer to the plant cell but lacks a transferable T-DNA. The T-DNA was incorporated onto a much smaller plasmid that can be engineered in *E. coli* and then transferred to *Agrobacterium*, where the *vir* genes mobilize the transfer of the T-DNA to plant cells. This binary vector system is commonly used to obtain transgenic plants. During *Agrobacterium* infection, the VirD2 excises the T-strand and covalently links it to tyrosine 29 (Tyr29) of VirD2. VirD2 guides the T-DNA through the Type IV secretion system (T4SS) into the plant cell, where the T-DNA integrates in the plant nucleus. VirD2 has nuclear localization signals (NLS) (Tinland et al., 1992; Tinland et al., 1995; Kralemann et al., 2022) and T4SS translocation signals at its C-terminus (Vergunst et al., 2000; Vergunst et al., 2005; Li and Christie, 2018).

*Agrobacterium* has been a useful tool to transform the nuclear genome of different eukaryotes, including plants (Chilton et al., 1977; Marton et al., 1979), yeast (Piers et al., 1996), algae (Kumar et al., 2004) and fungal cells (de Groot et al., 1998). *Agrobacterium* T-DNA delivery to chloroplasts would be highly desirable because direct transfer of transforming DNA to plastids in the female gametes would avoid the need for tissue culture and plant regeneration. Our goal is to reengineer the T-DNA transfer machinery to target DNA delivery to chloroplasts.

Important for our project was choosing an assay that yields unambiguous data about the localization of the reengineered VirD2 protein. Fluorescent proteins have been used to study the subcellular localization of *Agrobacterium* effector proteins. In one approach, a photostable variant of phototropin Light, Oxygen and Voltage (LOV) domain, phiLOV2.1, was employed. phiLOV2.1 was fused at the N-terminus of the VirE2, VirE3, VirD2, VirD5 and VirF effector proteins and subcellular localization of the effector proteins was determined by phiLOV2.1 fluorescence (Roushan et al., 2018). In the second approach, GFP fluorescence was used for subcellular localization. Because GFP-tagged effector proteins do not pass through the T4SS, movement of VirE2 was tracked by fusing it with part of the GFP fluorescent protein, GFP_11_, and complementing GFP_1-10_, in the plant cell (Li et al., 2020). The split GFP assay (Cabantous et al., 2005) that we used for subcellular localization of VirD2 is discussed in more detail below.

We report here successful targeting of VirD2 to plastids based on a split GFP assay where a VirD2 fused with GFP_11_ complements GFP_1-10_ in chloroplasts. Reengineering VirD2 will enable organellar genome engineering, extending the utility of *Agrobacterium* for DNA delivery beyond the nucleus.

## Results

### Experimental design

When a protein is targeted to the nucleus, all targeted molecules are concentrated at the target site. Detecting imported VirD2 in chloroplasts is more challenging than detection in the nucleus because there are about 100 chloroplasts in a tobacco leaf cell (Greiner et al., 2020), and the fluorescence signal is split 100 ways. To avoid ambiguities due to the low amount of VirD2 exported from *Agrobacterium* to chloroplasts, we decided to express the reengineered VirD2 from the plant nuclear genome (Figure 1A). For the assay, GFP is split into two parts (Cabantous et al., 2005). The first 10 β-strands of GFP (GFP_1-10_) are targeted to chloroplasts. The complementing small, 11^th^ ý-strand of the GFP (GFP_11_) is fused with chloroplast targeted VirD2, as discussed above. GFP_1-10_ and VirD2-GFP_11_ do not fluoresce on their own. Fluorescence can be detected only when VirD2-GFP_11_ is imported into chloroplasts where it complements GFP_1-10._ In our test system, both genes were expressed from a plant nuclear gene, therefore both GFP_1-10_ and VirD2-GFP_11_ were fused with a plastid-targeting transit peptide, that is cleaved off by the stromal processing protease after import (Sjuts et al., 2017) (Figure 1A).

**Figure 1.**
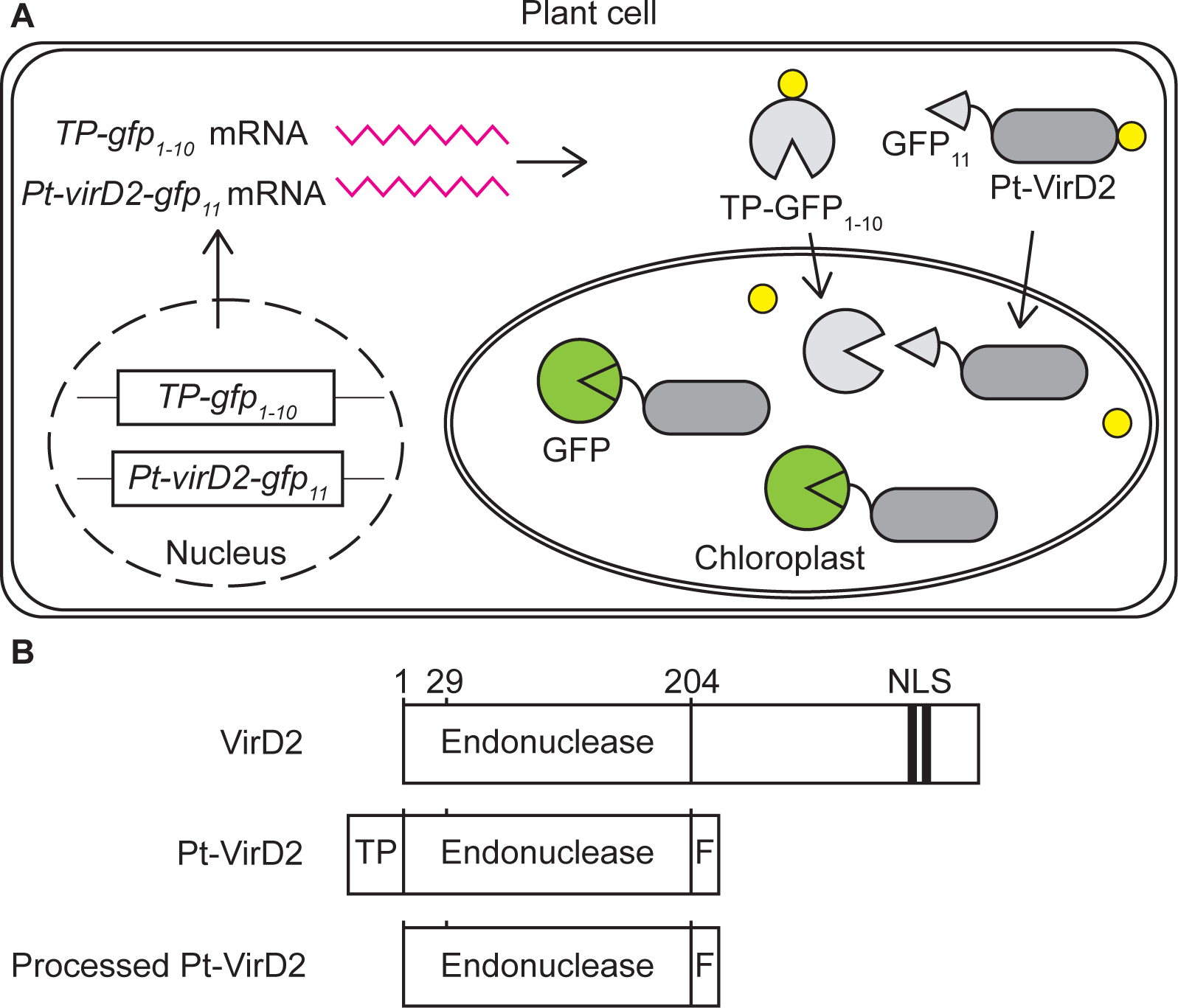
Schematic diagram of split GFP assay to confirm chloroplast targeting of VirD2. **A**, The plant cell, with plastid-targeted *TP-gfp_1-10_* and *Pt-virD2-gfp_11_* genes in the nucleus, and the TP-GFP_1-10_ and Pt-VirD2-GFP_11_ proteins in the cytoplasm and chloroplasts. Note that TP-GFP_1-10_ and Pt-VirD2-GFP_11_ complement creating a fluorescent protein and that the transit peptide (TP) symbolized by a yellow circle is present in the cytoplasm, and it is cleaved off after import in the chloroplast. **B**, Schematic structure of VirD2, Pt-VirD2 and processed Pt-VirD2 (TP removed) proteins. For further explanation see text.

When designing the plastid *virD2* gene (*Pt-virD2*), we relied on information dissecting VirD2 function. AS starting point, we used the FLAG-VirD2-204-F construct described by van Kregten et al. (van Kregten et al., 2009) where we replaced the FLAG tag at the N-terminus of with the Rubisco small subunit (SSU) transit peptide (TP) (Gnanasambandam et al., 2007), (Figure 1b). In *Pt-VirD2* the first 204 amino acids encode the endonuclease domain required for excision of the T-strand at the T-DNA borders and linkage of the T-strand to Tyr29 of VirD2. The C-terminal end of *Pt-virD2* encodes the T4SS signal.

### Split GFP complementation assay in *E. coli*

Placement of the GFP_11_ peptide at the N- or C-terminus of Pt-VirD2 and inclusion of linkers between GFP_11_ and the adjacent protein sequence are critical for complementation (Cabantous et al., 2005). To obtain information about the feasibility of complementation, we first tested Pt-VirD2 complementation in *E. coli* (Figure 2).

**Figure 2.**
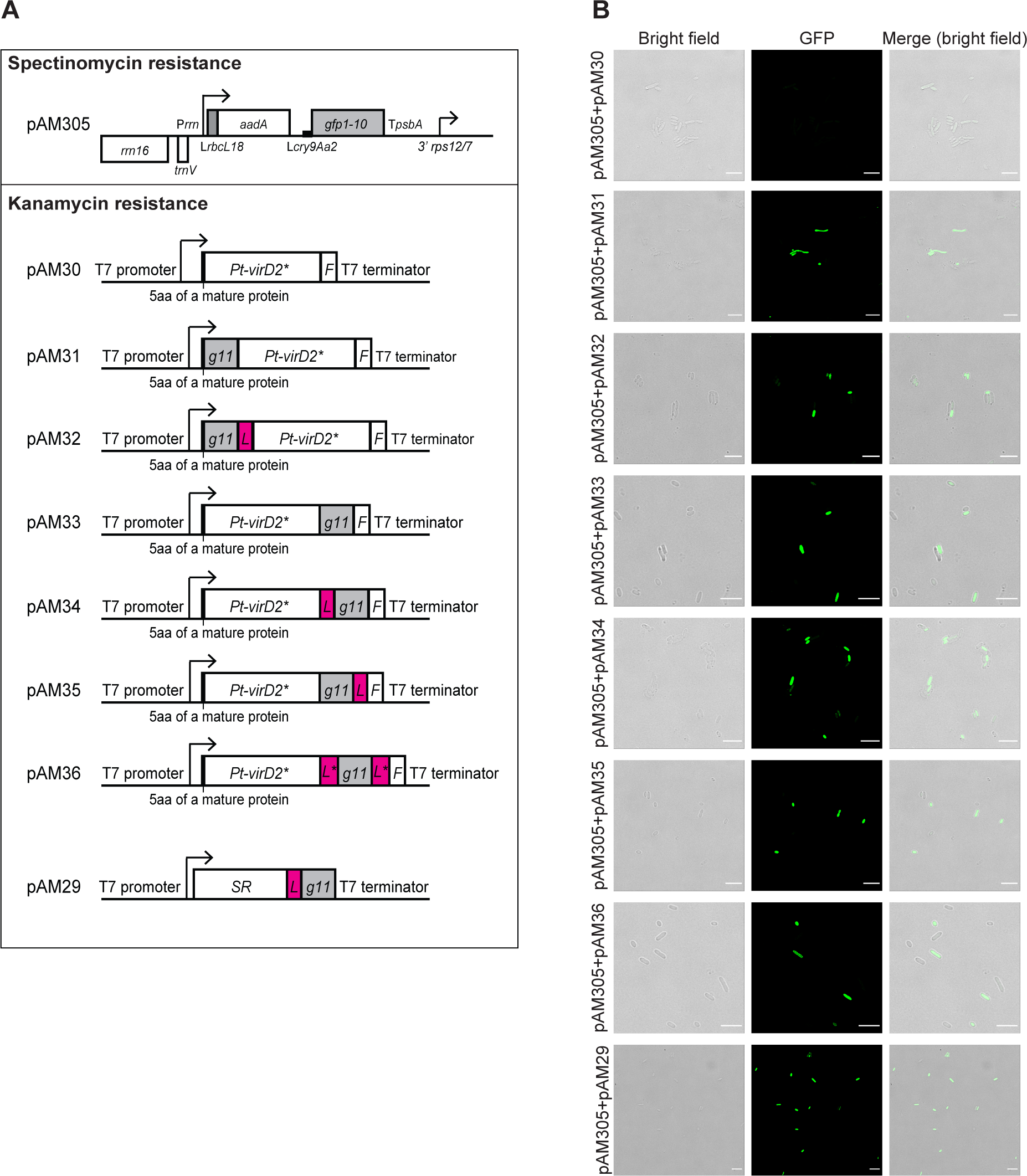
VirD2-GFP_11_ and GFP_1-10_ complementation in *E. coli* grown at 28 °C. **A,** Expression of GFP_1-10_ in plasmid pAM305 and Pt-VirD2 derivatives in pAM30-pAM36. pAM305 carries a spectinomycin resistance (*aadA*) marker gene in a dicistronic chloroplast vector, a pMRR13 derivative (Yu et al., 2020). The Pt-VirD2 derivatives are expressed from the pET28a vector carrying a kanamycin resistance marker. Sulfite reductase (SR) in pAM29 is the positive control, expressing SR-linker (L) – GFP_11_ fusion (Cabantous et al., 2005). The linkers were L, (GGGS)x2 or L* (GGGS). F is the T4SS signal of VirF (F). **B**, Fluorescence in *E. coli* confirms complementation of GFP_1-10_ and Pt-VirD2-GFP_11_ proteins. The bar is 5 μm. Data were collected on a Leica Stellaris8 confocal microscope.

GFP_1-10_ was expressed from a pUC plasmid carrying a spectinomycin resistance marker (pAM305) and VirD2-GFP_11_ variants in a pET28a vector carrying a kanamycin resistance marker. Selection for spectinomycin resistance and kanamycin resistance ensured that both plasmids were present in *E. coli*. The VirD2-GFP_11_ plasmids expressed a variant lacking the TP but had the five amino acids of mature SSU at the N-terminus, the form that will be present when the TP is cleaved off after import into chloroplasts. Some of the VirD2 variants carried the GFP_11_ region at the N-terminus, others between the truncated VirD2 C-terminus and the VirF T4SS signal. In some of the plasmids the 16 amino acid GFP_11_ peptide is flanked by linker sequences (Figure 2A). The positive control for GFP complementation was plasmid pAM29 expressing sulfite reductase-GFP_11_ fusion protein, that proved to be useful to develop the original split-GFP assay (Cabantous et al., 2005). The negative control was pAM30, Pt-VirD2 without GFP _11_ peptide. Each of the VirD2-GFP_11_ proteins complemented GFP_1-10_, as indicated by fluorescence under the confocal microscope when the *E. coli* were grown at 28°C (Figure 2B). However, complementation was absent when grown at the standard 37°C. We hypothesize that protein synthesis at the lower temperature was slower, allowing time for protein folding and assembly. The idea to test expression at lower temperatures came from previous VirD2 expression studies in *E. coli* (Pelczar et al., 2004; Matsuoka, 2015).

### Split GFP complementation assay in tobacco chloroplasts

Successful complementation of proteins in the *E. coli* assay set the stage for testing subcellular protein targeting in plants. First, we generated multiple, independently transformed tobacco plants which accumulate GFP_1-10_ at different levels dependent of the site of insertion in the nuclear genome (Figure 3). We then tested the GFP_1-10_ expressing lines for background fluorescence under the confocal microscope. We chose the line with the lowest GFP_1-10_ level (pAM407-10), in which no fluorescence was detectable in the GFP channel, as in the wild-type chloroplasts (Figure 4B).

**Figure 3.**
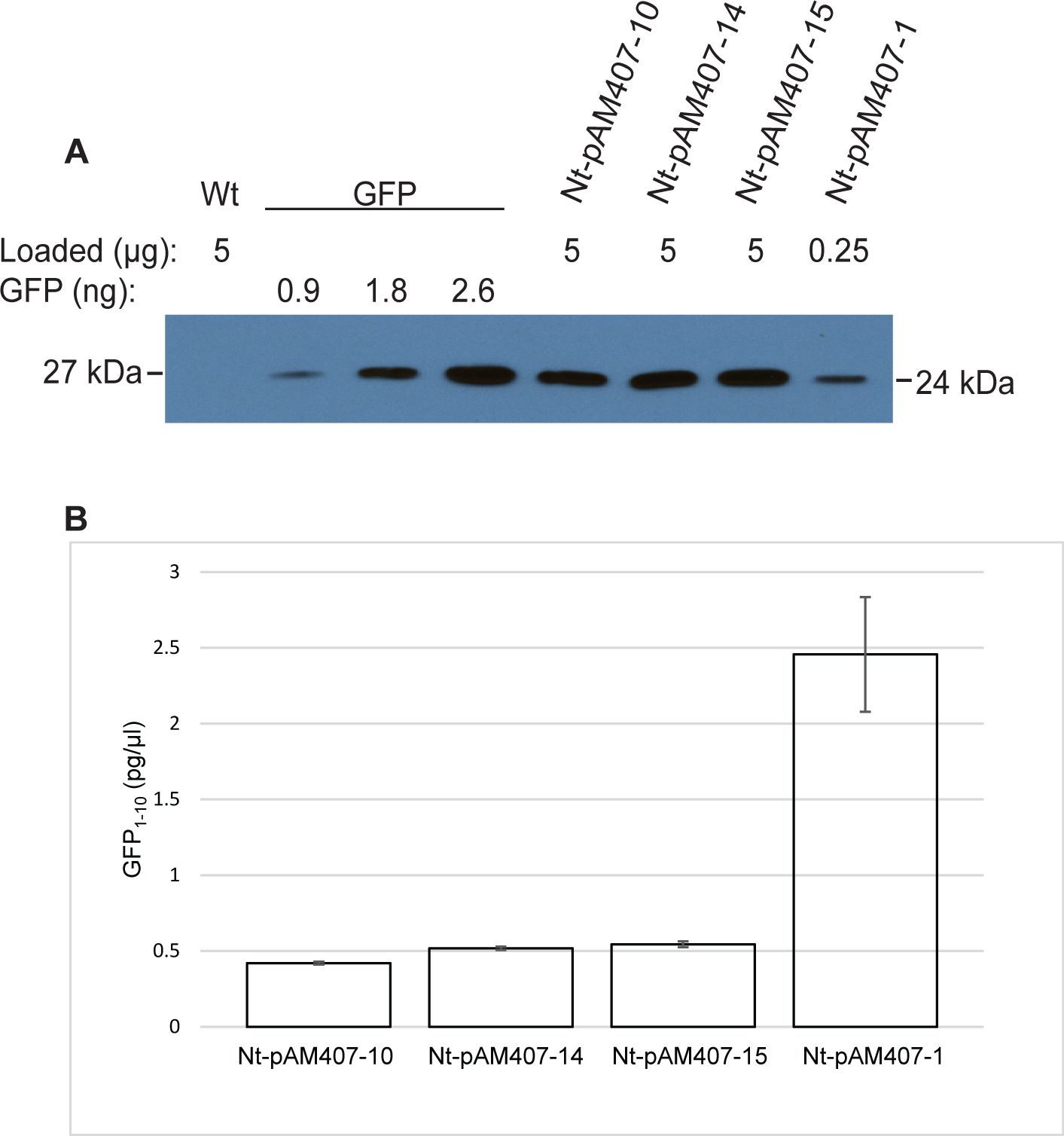
Quantification of GFP_1-10_ in the leaves of Nt-pAM407 tobacco plants. **A**, Representative western blot. 5 μg total soluble protein was loaded per lane, except the high expressor line Nt-pAM407-1 (0.25 μg per lane). Reference GFP protein was obtained by dilution of protein extracts from chloroplast expressed GFP. **B**, GFP_1-10_ concentration (pg/μl) in leaf protein extracts of individual Nt-pAM407 transgenic lines. The values are an average of three independent measurements.

**Figure 4.**
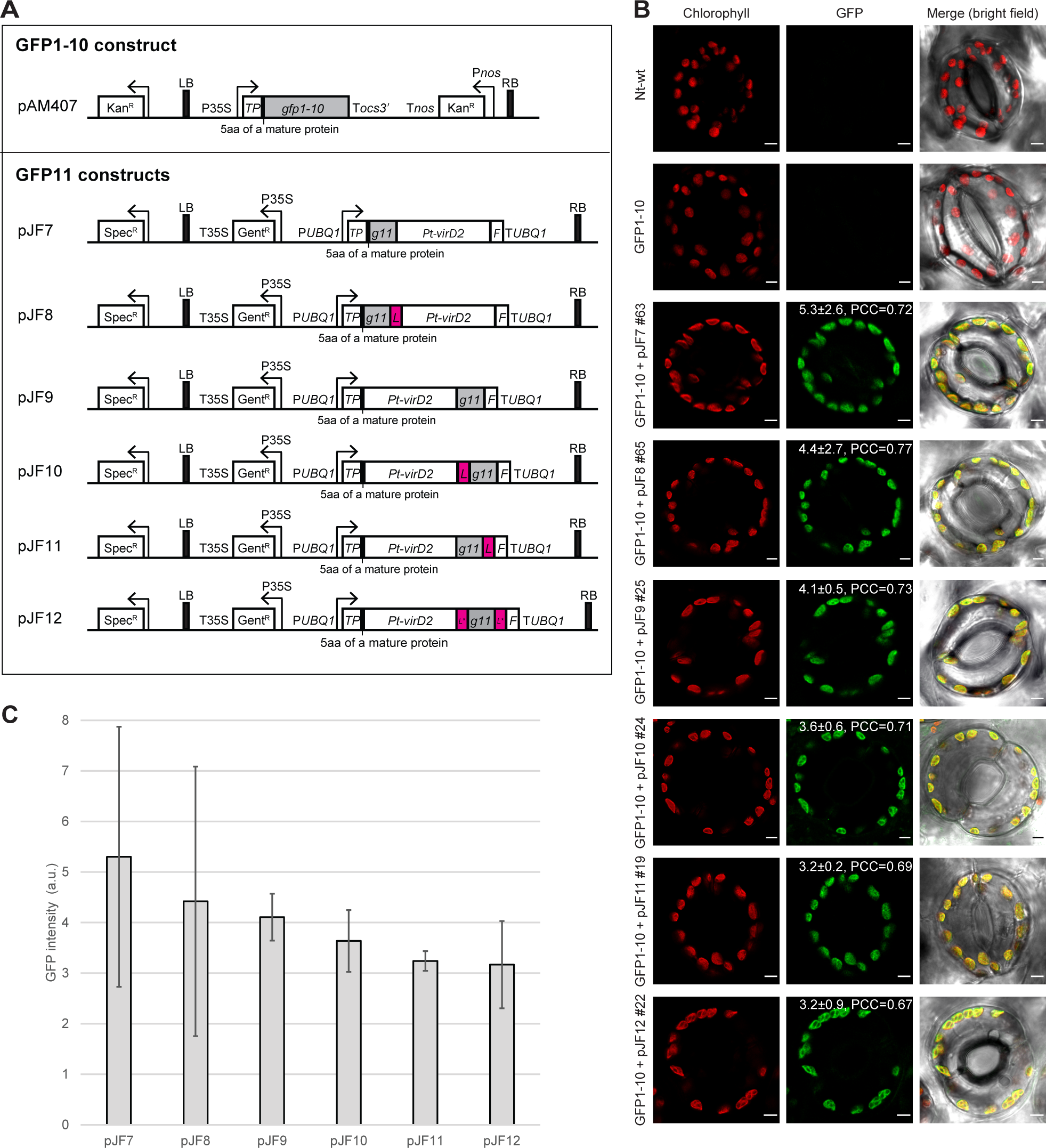
Chloroplast targeting of Pt-VirD2-GFP_11_ confirmed by complementation with GFP_1-10_ in chloroplasts. **A**, Binary vectors to test Pt-VirD2-GFP_11_ and GFP_1-10_ complementation in chloroplasts. Pt-VirD2 was fused with 16 amino acids of the GFP-11 peptide (RDHMBLHEYBNAAGIT) at the N- or C-terminus with or without a linker [L, (GGGS)x2; L*, GGGS], and the T4SS signal of VirF (F). **B**, Confocal images of GFP complementation in chloroplasts. The chlorophyll signal (excitation 633 nm; emission 648 to 700 nm) and GFP signal (excitation 488 nm; emission 498 to 548 nm) were visualized using a Leica TCS SP8 confocal microscope. Scale bar, 5 µm. Mean GFP intensity, standard deviation and Pearson’s correlation coefficient (PCC) were calculated using Fiji (ImageJ) software (Schindelin et al., 2012) and indicated on GFP panels. The values are an average of >100 chloroplasts each in five independently transformed lines. **C**, GFP intensity in complementing lines expressed in arbitrary units (a.u.).

To test complementation in plants, the selected GFP _1-10_ line (pAM407-10 line; kanamycin resistant) was transformed with the *Pt-virD2-gfp_11_* genes (pJF7-pJF12 binary vectors; gentamycin resistant; Figure 4A). The *Pt-virD2-gfp_11_* genes are expressed from a Ubiquitin 1 gene promoter/terminator (P*UBQ1*/T*UBQ1*) cassette where they are transcribed, then translated in the cytoplasm and imported into chloroplasts (Figure 1A). Figure 4B shows red fluorescence of chlorophyll, green fluorescence when Pt-VirD2-GFP_11_ is expressed at a high level to give a strong signal after assembly with GFP_1-10_, and colocalization of the two in stoma guard cells in an image taken under bright field microscopy (Figure 4B). Although the *Pt-virD2-GFP_11_* genes were expressed from the same promoter-terminator cassette, only some of the independently transformed lines produced Pt-VirD2-GFP_11_ at high-enough level to yield complementation (Table 1). In the wild-type control (no *gfp* gene) and pAM407-10, expressing only *gfp_1-10_*, no fluorescence could be detected in the GFP channel. Fluorescence intensity in chloroplasts expressing GFP1-10 (pAM407 construct) and one of the six VirD2-GFP_11_ fusion proteins was in the range of 3.2 to 5.3. Fluorescence of GFP_11_ N-terminal fusions with VirD2 appear to be higher. However, these values are not significantly different (Figure 4C). The Pearson’s correlation coefficients (PCC) of chlorophyll and GFP signals indicate strong colocalization (Zinchuk et al., 2013).

**Table 1.** Complementation in tobacco by transforming GFP_1-10_ (Nt-pAM407-10, kanamycin resistant) line with VirD2-GFP_11_ vectors (pJF7-pJF12, gentamycin resistant). We listed the number of double-(kanamycin, gentamycin) resistant lines and the number with detectable GFP complementation.

| VirD2-GFP<sub>11</sub> vectors | Kan/Gent resi. | + GFP |
| --- | --- | --- |
| pJF7 | 41 | 11 |
| pJF8 | 53 | 11 |
| pJF9 | 36 | 13 |
| pJF10 | 46 | 14 |
| pJF11 | 42 | 12 |
| pJF12 | 36 | 10 |

## Discussion

Our goal is transforming plastids using *Agrobacterium* for T-DNA delivery. We previously demonstrated that a protein can be targeted from *Agrobacterium* to chloroplasts using the phiC31 phage site-specific integrase (Int). Int was engineered to have the VirF T4SS signal at the C-terminus for export from *Agrobacterium* to the plant cell and the Rubisco small subunit transit peptide (SSU-TP) for targeting to chloroplasts. Int visitation was detected by excision of a marker gene (Matsuoka and Maliga, 2021). We report here that the second step needed to convert *Agrobacterium* for T-DNA delivery to plastids, reengineering VirD2 itself, was achieved by truncating the coding region to remove the C-terminal NLSs and then adding a T4SS signal at the C-terminus and SSU-TP at the N-terminus.

So far VirD2 has always been targeted to the nucleus. We show the role of VirD2 in nuclear gene transformation on Figure 5A. Reengineering VirD2 for chloroplast targeting is a major advance, opening the way for DNA delivery to chloroplasts. For this, Pt-VirD2 will be cloned in the binary vector between T-DNA border sequences (Figure 5). For successful T-DNA transfer, it is important that VirD2 maintains endonuclease activity when fused with the tobacco Rubisco SSU-TP (Matsuoka, 2015; Roushan et al., 2018). The virulence plasmid will lack the wild-type VirD2 to avoid competition between VirD2 and Pt-VirD2 in *Agrobacterium*. An *Agrobacterium* strain lacking the wild type *virD2* gene is available (Li et al., 2020).

**Figure 5.**
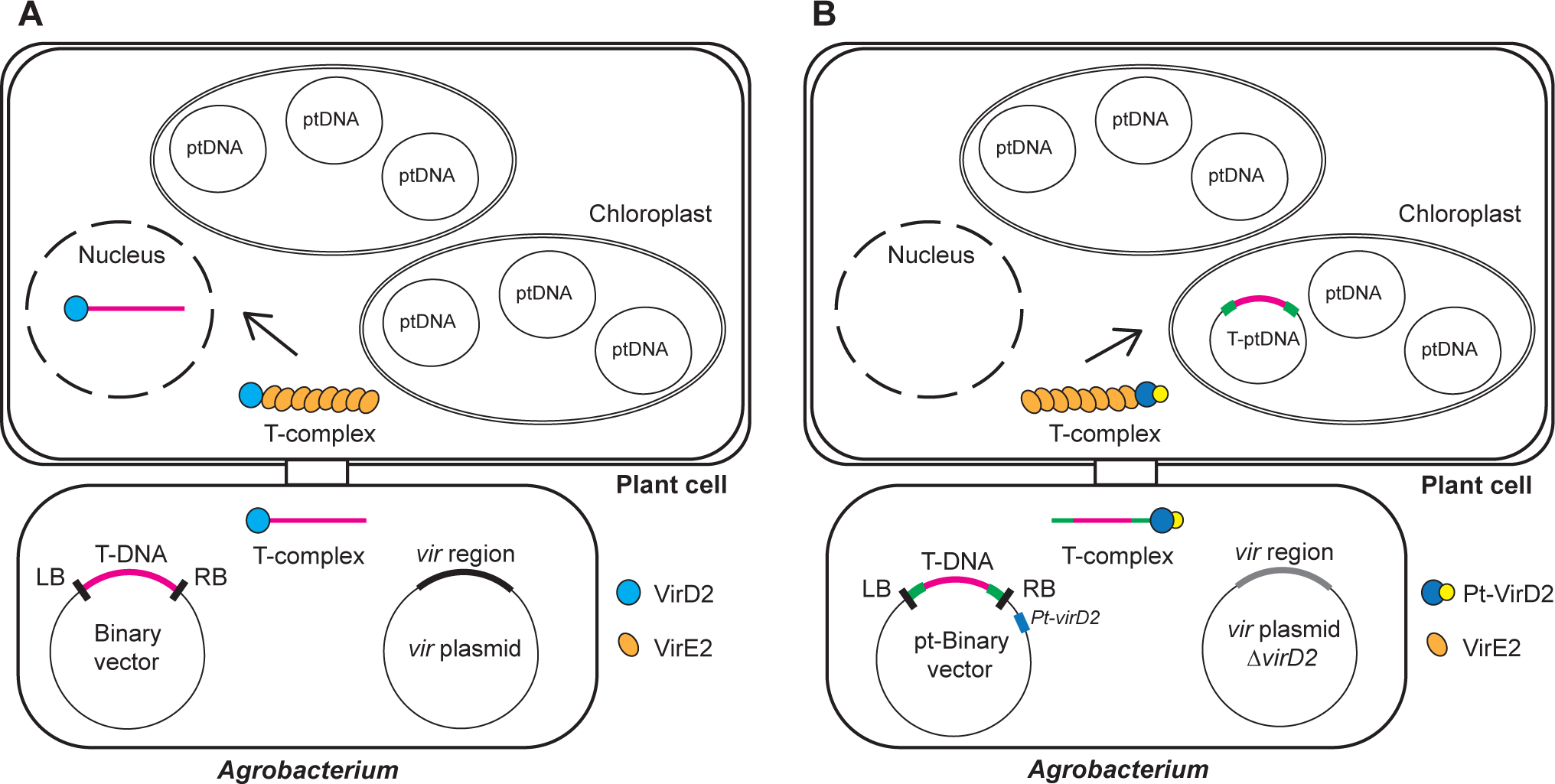
Conceptual design of chloroplast transformation by *Agrobacterium*. **A**, *Agrobacterium* transformation of the nucleus. **B**, *Agrobacterium* transformation of chloroplasts with the engineered Pt-VirD2. Chloroplast targeting region of the Pt binary vector is in green.

Reengineering VirE2, the single-stranded DNA binding protein that protects the T-strand from degradation may not be necessary because VirE2 NLSs do not function in plant cells (Li et al., 2020). It appears that the T-complex is transported into the host nucleus in a cooperative manner, where VirD2 provides guidance on the “head” through its NLS-mediated interaction with plant importin α, whereas VirE2 provides the assistance on the lateral side of the T-complex via VirE2 and host nucleoporin CG1 interaction (Li et al., 2020). If VirD2 is targeted to plastids, VirE2 is likely to follow. Because the transforming DNA integrates in the plastid genome by homologous recombination via the plastid targeting sequences (Maliga, 2004), it is sufficient to provide plastid sequences for targeted insertion of transgenes in the plastid genome (Figure 5B). This makes engineering effector proteins involved in T-DNA integration, such as VirF, unnecessary.

Once *Agrobacterium*-mediated DNA delivery yields stable transplastomic plants in tobacco leaf mesophyll cells, it will be time to attempt plastid transformation in Arabidopsis. Although the female gamete contains multiple plastids, germinating the seed on selective spectinomycin medium will favor retention of transplastomic plastids. Recovery of rare transplastomic plants in the seed progeny after a floral dip in an *Agrobacterium* suspension will be similar to recovering rare paternal plastids in the seed progeny (Ruf et al., 2007; Svab and Maliga, 2007). Plastid transformation by the floral dip protocol will avoid tissue culture, making plastid transformation readily availably for the research community.

## Materials and methods

### Plasmids for the split GFP complementation assay

Plastid vector pAM305 is a pMRR15 derivative (Yu et al., 2020) in which *gfp* was replaced with *gfp_1-10_*. The pAM305 DNA sequence is included in Table S1.

To test VirD2 complementation in *E. coli*, plastid *virD2* (*Pt-virD2*\*) genes were constructed to yield processed forms of proteins in *E. coli*. The processed forms are present after import into chloroplasts, lacking the Rubisco small subunit transit peptide other than the 5 amino acids (MQVWP) left after cleavage by the stromal processing protease. VirD2 was fused with 16 amino acids of the GFP-11 peptide (RDHMBLHEYBNAAGIT) at the N- or C-terminus with or without a linker [L, (GGGS)x2; L*, GGGS], and the T4SS signal of VirF (F). The seven processed Pt-VirD2 genes (pAM30-pAM36; Figure 2A) are expressed from a pET28a vector carrying a kanamycin resistance marker. pAM29 encodes a sulfite reductase (SR) with a C-terminal 2xLinker and GFP11 fusion (Cabantous et al., 2005).

pAM407 is a pBIN19 binary plasmid derivative (Bevan, 1984) for the expression of *gfp_1-10_* in a cauliflower mosaic virus 35S promoter/terminator (P35S/T35S) cassette. The plant marker is kanamycin resistance. The pAM407 DNA sequence is given in Table S1. The nuclear encoded, chloroplast-targeted *Pt-virD2-gfp_11_* genes were introduced into the tobacco nucleus using a gentamycin resistant pPZP221 binary vector (Hajdukiewicz et al., 1994) (pJF7-pJF12). The *Pt-virD2-gfp_11_* genes were expressed from a Ubiquitin 1 promoter/terminator cassette (Liu et al., 2015).

### GFP complementation in *E. coli*

*E. coli* strain BL21 (DE3) was transformed with pAM305 and pAM29-36 by heat-shock transformation. Transformants were selected on LB containing 100 mg/L spectinomycin and 50 mg/l kanamycin and incubated at 37°C overnight. Transformants were then transferred onto LB plates containing the same antibiotics and 1 mM IPTG and were incubated at 28°C overnight. GFP complementation was detected by fluorescence using a Leica Stellaris 8 microscope. The excitation/emission spectra for GFP are 488 nm/510 to 520 nm.

### Tobacco leaf transformation with chloroplast targeted GFP_1-10_ and VirD2-GFP_11_

*Nicotiana tabacum cv.* Petit Havana leaf discs were transformed with *Agrobacterium* EH105 (Hood et al., 1993) carrying vector pAM407. The *Agrobacterium* was grown in LB medium containing 25 mg/L chloramphenicol and 50 mg/L kanamycin. Tobacco leaf discs were dipped briefly in an *Agrobacterium* suspension and then grown on RMOP medium in the absence of selection. After two days the leaf disks were transferred to a selective RMOP shoot regeneration medium containing 500 mg/L carbenicillin and 50 mg/L kanamycin (Hajdukiewicz et al., 1994). The regenerated shoots were rooted on RM medium containing 500 mg/L carbenicillin and 50 mg/L kanamycin and then transferred to the greenhouse. GFP_1-10_ expression level in the leaves was determined by western blot analyses.

The *Pt-virD2-gfp_11_* genes in vectors pJF7-pJF12 were also introduced into the nucleus of *N. tabacum* expressing GFP_1-10_ by cocultivation with EHA105. Because the binary vectors pPZP221 carry spectinomycin resistance as bacterial marker, *Agrobacterium* was grown in LB medium containing 25 mg/L chloramphenicol and 100 mg/L spectinomycin. Selection of transgenic events was carried out on RMOP medium containing 500 mg/L carbenicillin and 100 mg/L gentamicin. Shoots were then rooted on RM medium containing 500 mg/L carbenicillin and 100 mg/L gentamicin.

### Extraction and quantification of GFP_1-10_

About 100 mg of ground leaf powder was resuspended in 200 µl of protein extraction buffer containing 50 mM HEPES (pH7.5), 1 mM EDTA, 10 mM potassium acetate, 5 mM magnesium acetate, 10 mM DTT, 2 mM PMSF, protease inhibitor cocktail (15μl/l; Sigma-Aldrich). Samples were vortexed for 20 sec and then placed on ice for 10 min. The supernatants were collected after centrifugation at 14,000xg for 5 min at 4°C. The concentration of total soluble protein was determined using the Bradford reagent (Bio-Rad) with bovine serum albumin as standard. For western blot, 5 µg of total soluble protein was loaded per lane, except the high expressor Nt-pAM407-1 line (0.25 µg per lane). GFP_1-10_ were detected using Living Colors® A.v. Monoclonal Antibody (JL-8) (632381; Clontech) in 1:2,000 dilution, anti-Mouse IgG (whole molecule)– Peroxidase antibody produced in goat (A4416; Sigma-Aldrich) in 1:18,000 dilution, and then Clarity Western ECL Substrate (Bio-Rad). GFP (0.9, 1.8 and 2.6 ng) from a chloroplast expressed GFP plant was used as standards for GFP_1-10_ quantification. The signal intensity was calculated with Fiji (ImageJ) software (Schindelin et al., 2012) using three different experimental replicates.

### Confocal microscopy and image processing

Leaf discs of Nt-pAM407-10 transformed with the pJF vectors were analyzed by a Leica TCS SP8 microscope. The chlorophyll signal (excitation 633 nm; emission 648 to 700 nm) and GFP signal (excitation 488 nm; emission 498 to 548 nm) were visualized using a Leica TCS SP8 confocal microscope. Mean intensity, standard deviation, and Pearson’s correlation coefficient (PCC) were calculated using Fiji (ImageJ) software (Schindelin et al., 2012) and are indicated on GFP panels in Figure 4B). The values are an average of at last 100 chloroplast reads in five independently transformed lines per construct.

## Supplemental Data

The following materials are available in the online version of this article.

**Supplemental Table S1**. DNA sequence of plasmids used for complementation in the split GFP assay in *E. coli* and in tobacco chloroplasts.

## Acknowledgements

We thank Dr. Dongmeng Li, Rutgers University, for her advice on confocal microscopy and the Waksman Institute Shared Imaging Facility. This work was supported by National Science Foundation grants MCB 1716102, IOS 2037155 and IOS 2224861 to P.M. and Kerry A. Lutz.

## Author contributions

P.M. and A.M. designed the experiments. A.M. and J.L.F. performed the experiments, P.M. and A.M. wrote the article.

## Conflict of interest

The authors declare that they have no conflict of interest.

## Data availability statement

The nucleotide sequences are included in Table S1. The author responsible for distribution of materials integral to the findings presented in this article in accordance with the policy described in the Instructions for Authors (https://academic.oup.com/plphys/pages/General-Instructions) is Pal Maliga. Biological Materials will be made available pending on the execution of a Biological Materials Transfer Agreement with Rutgers University.

---

**Supplemental Table S1.**
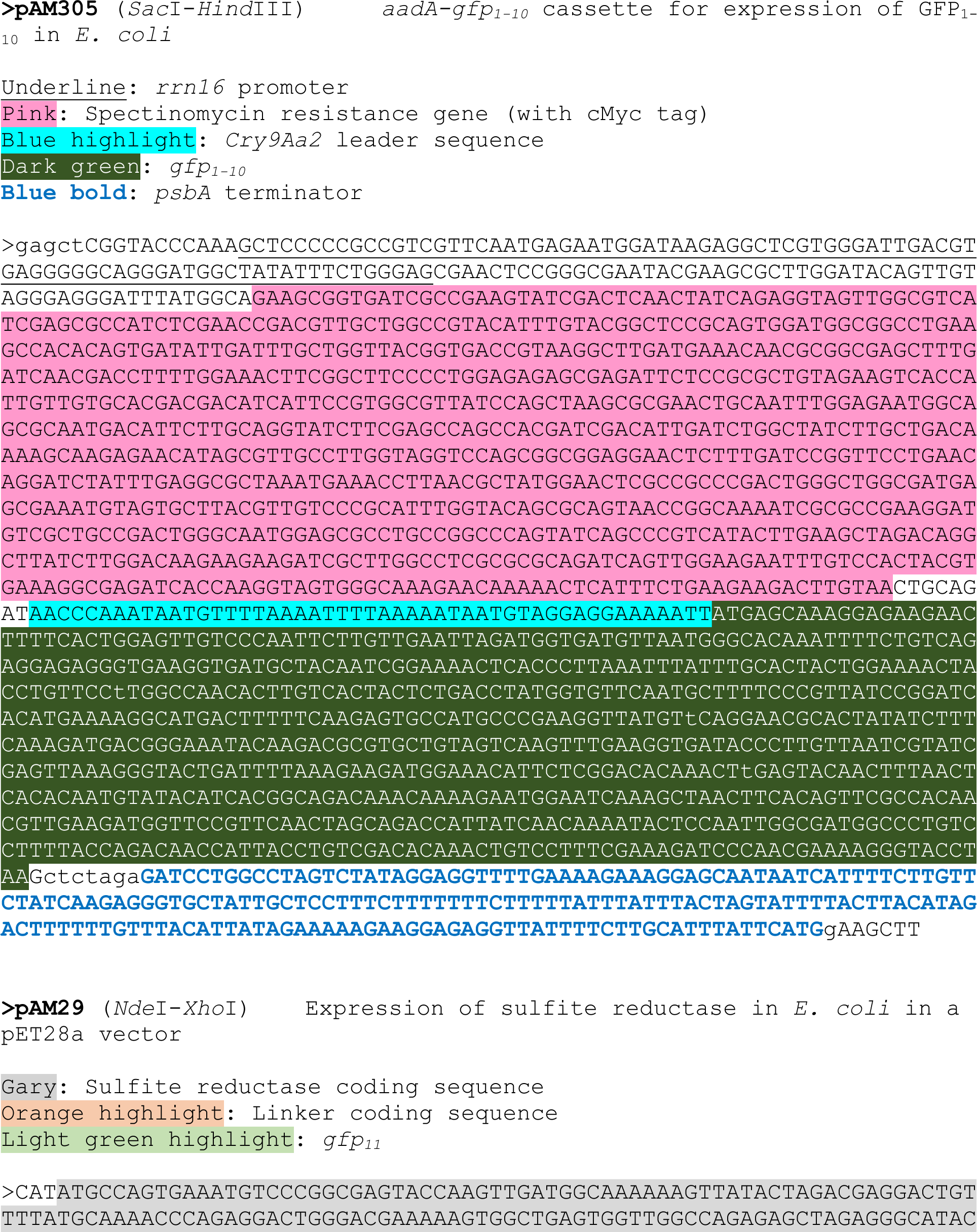
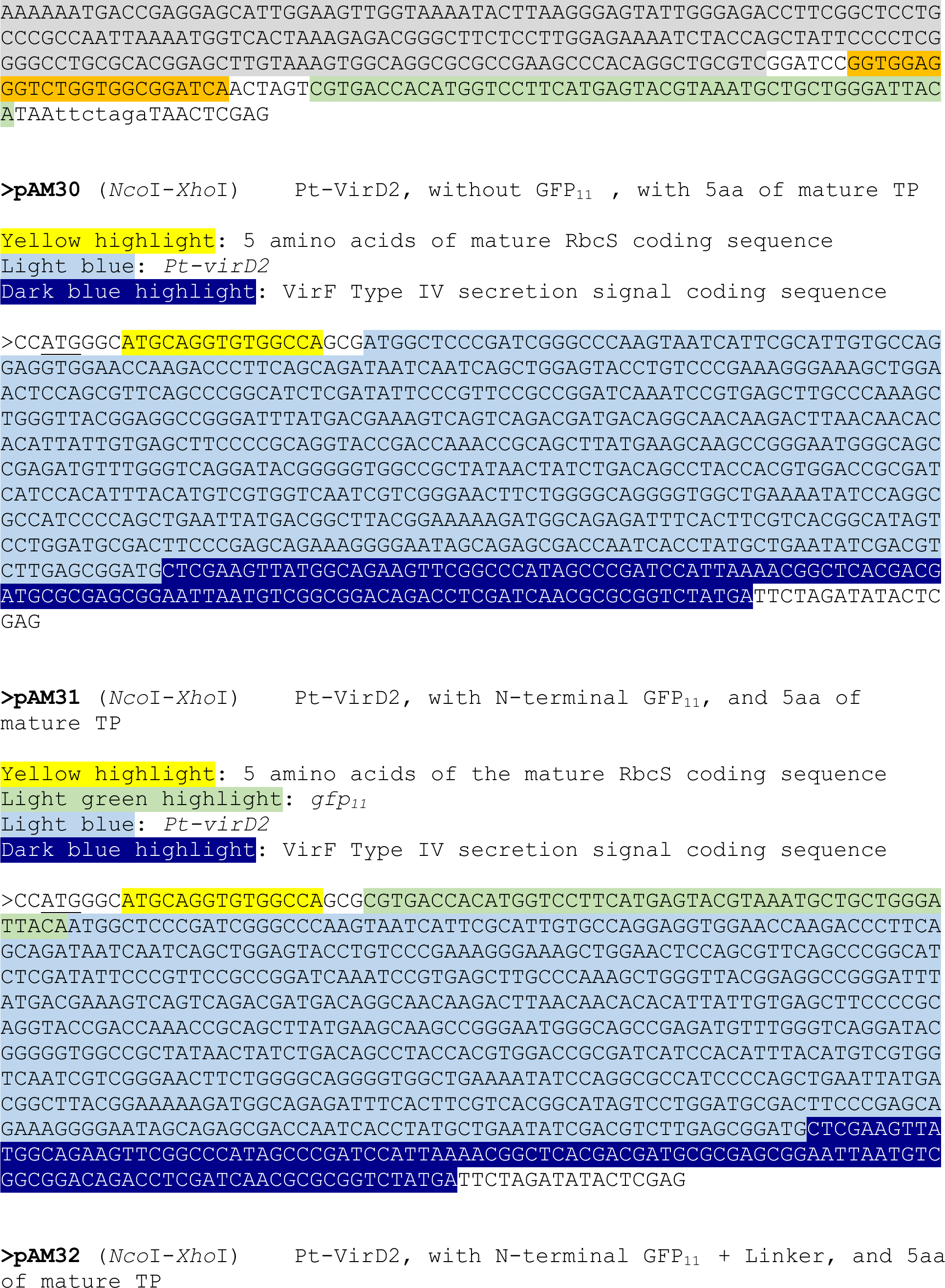
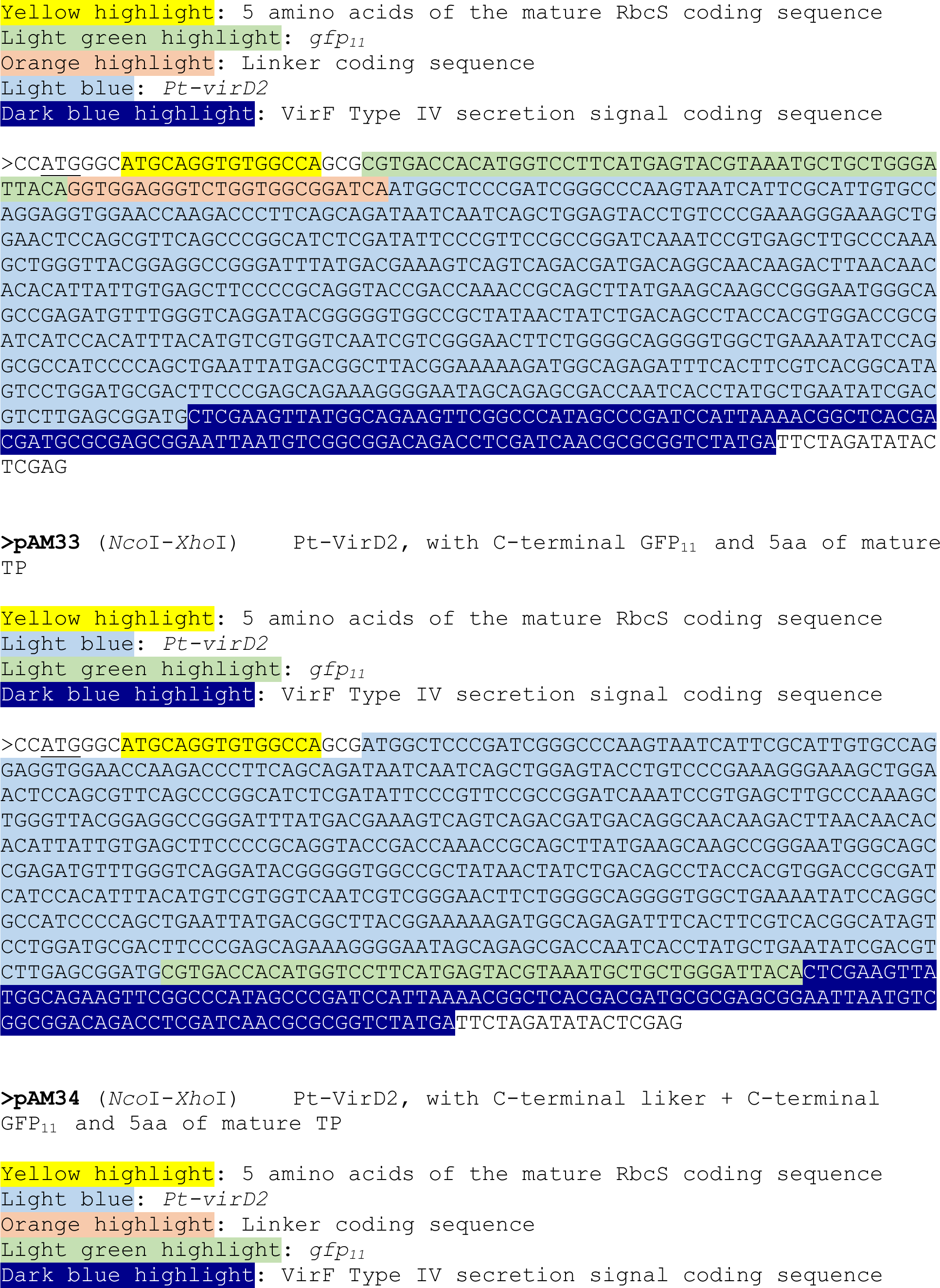
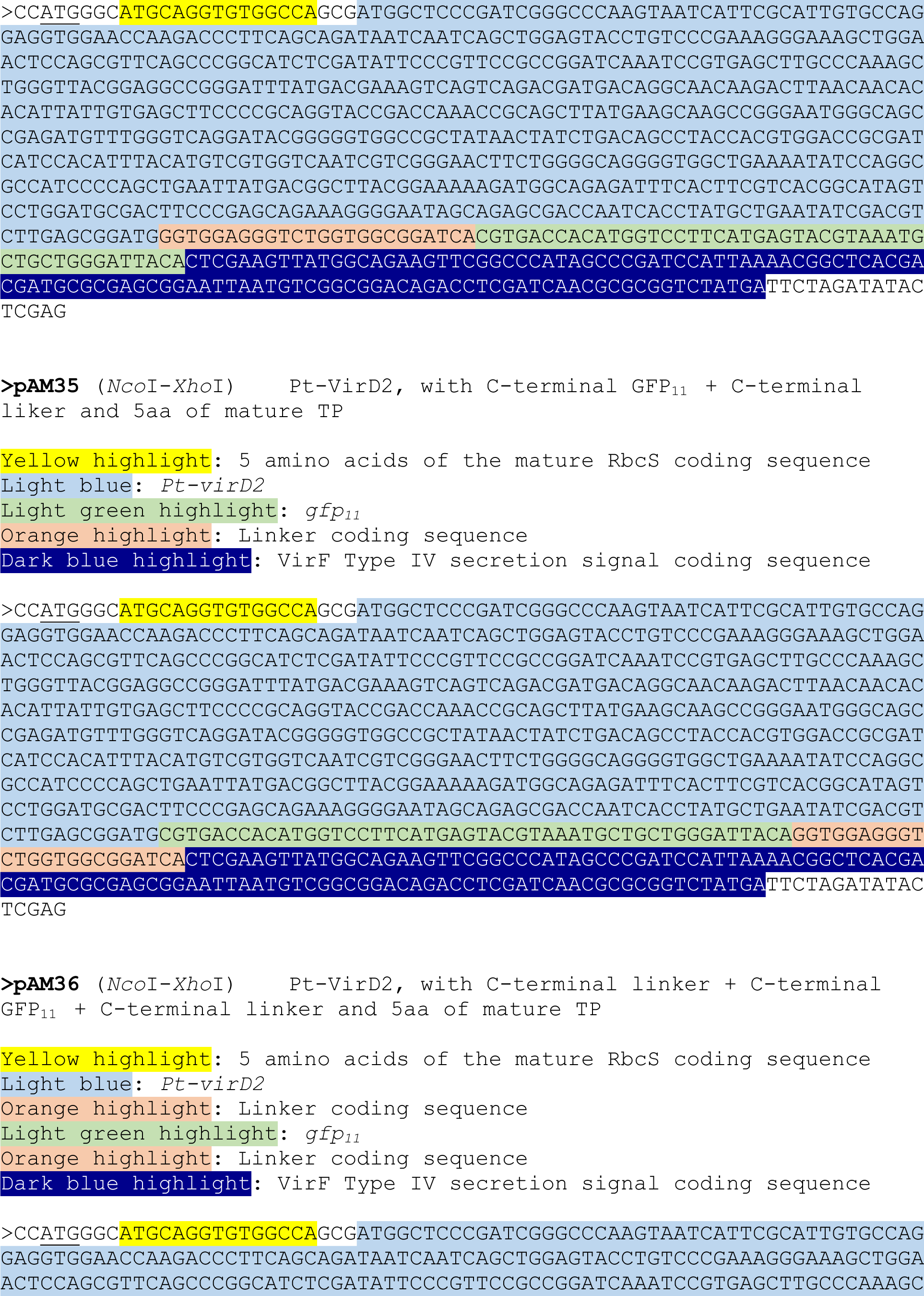
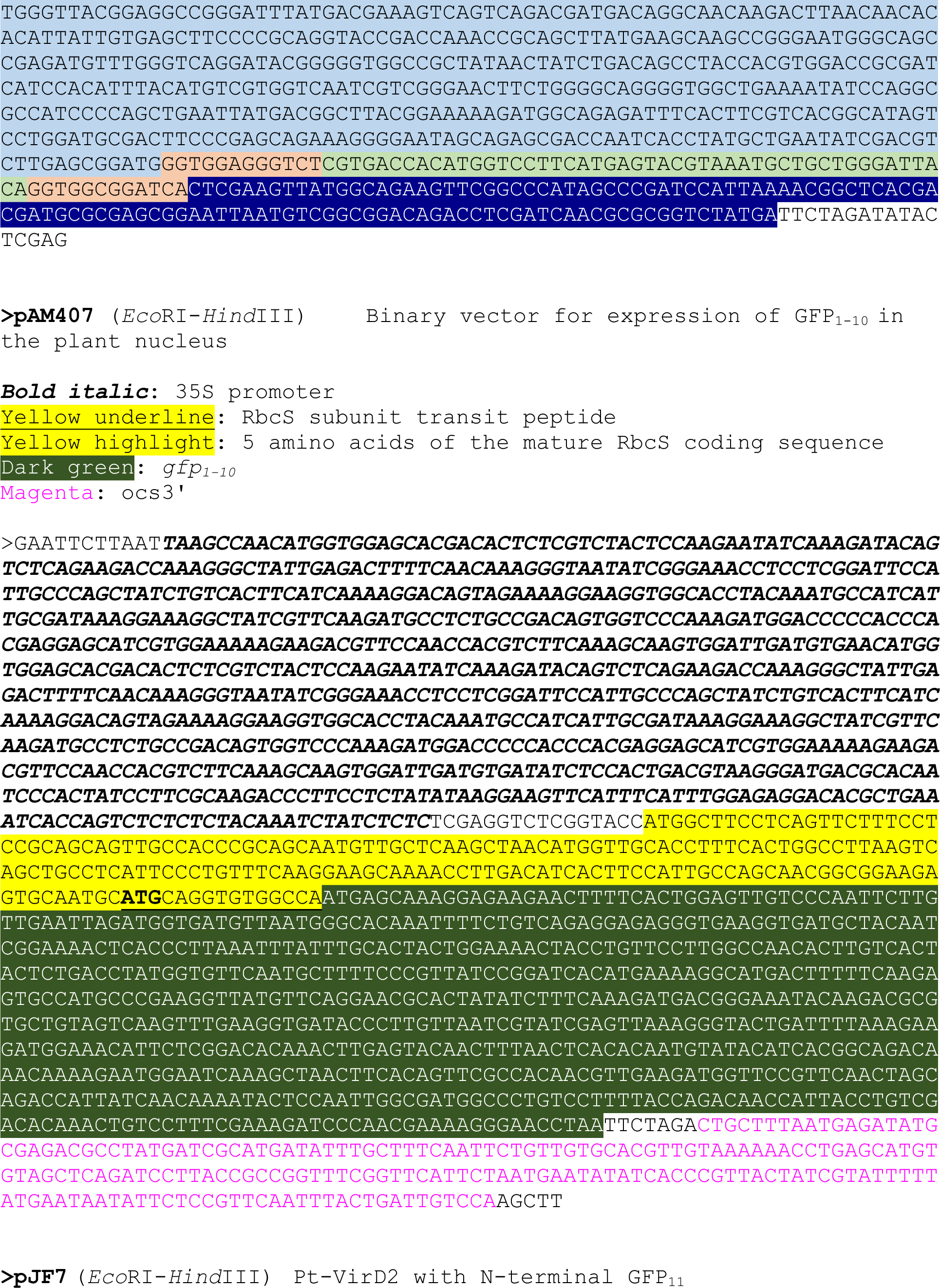
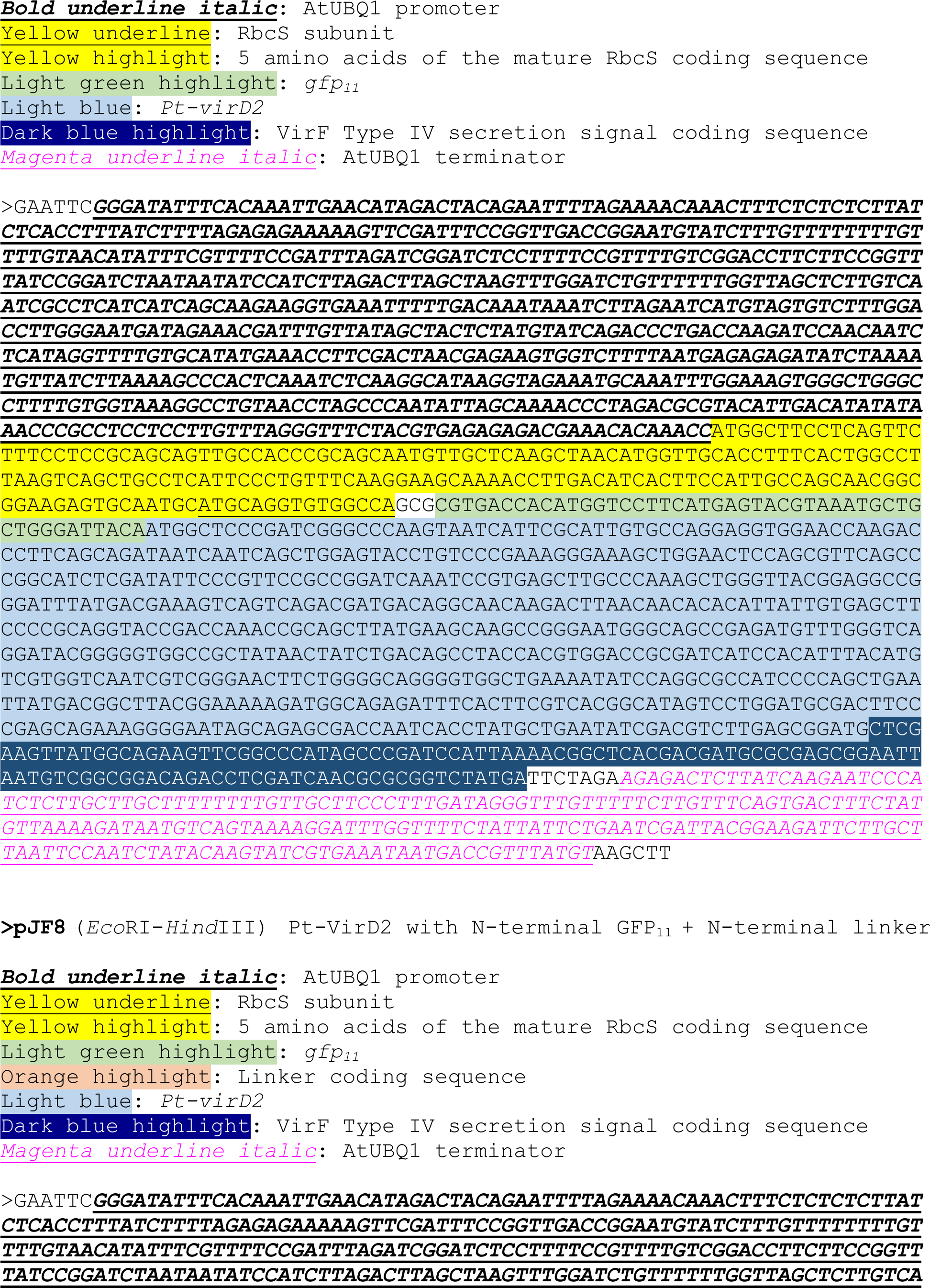
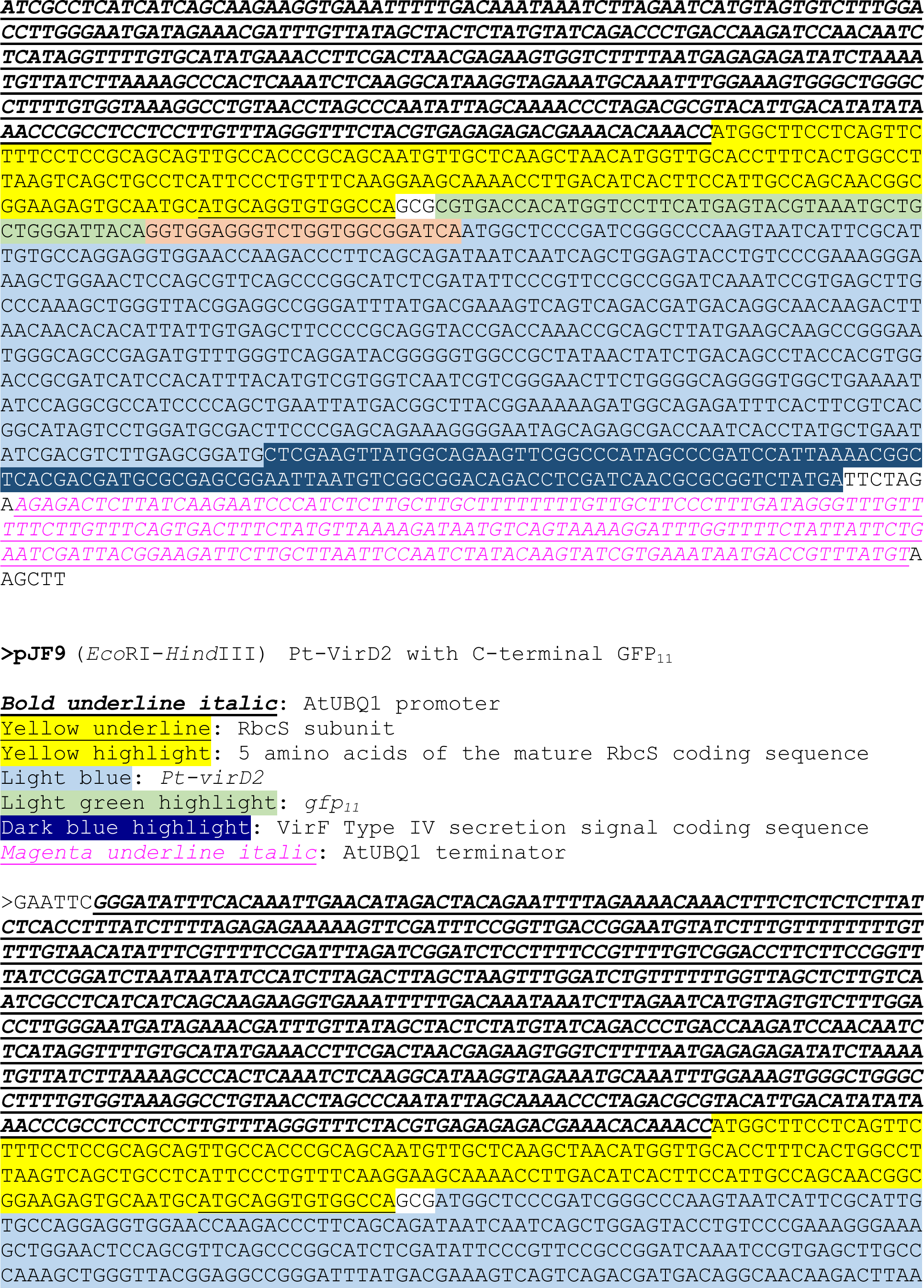
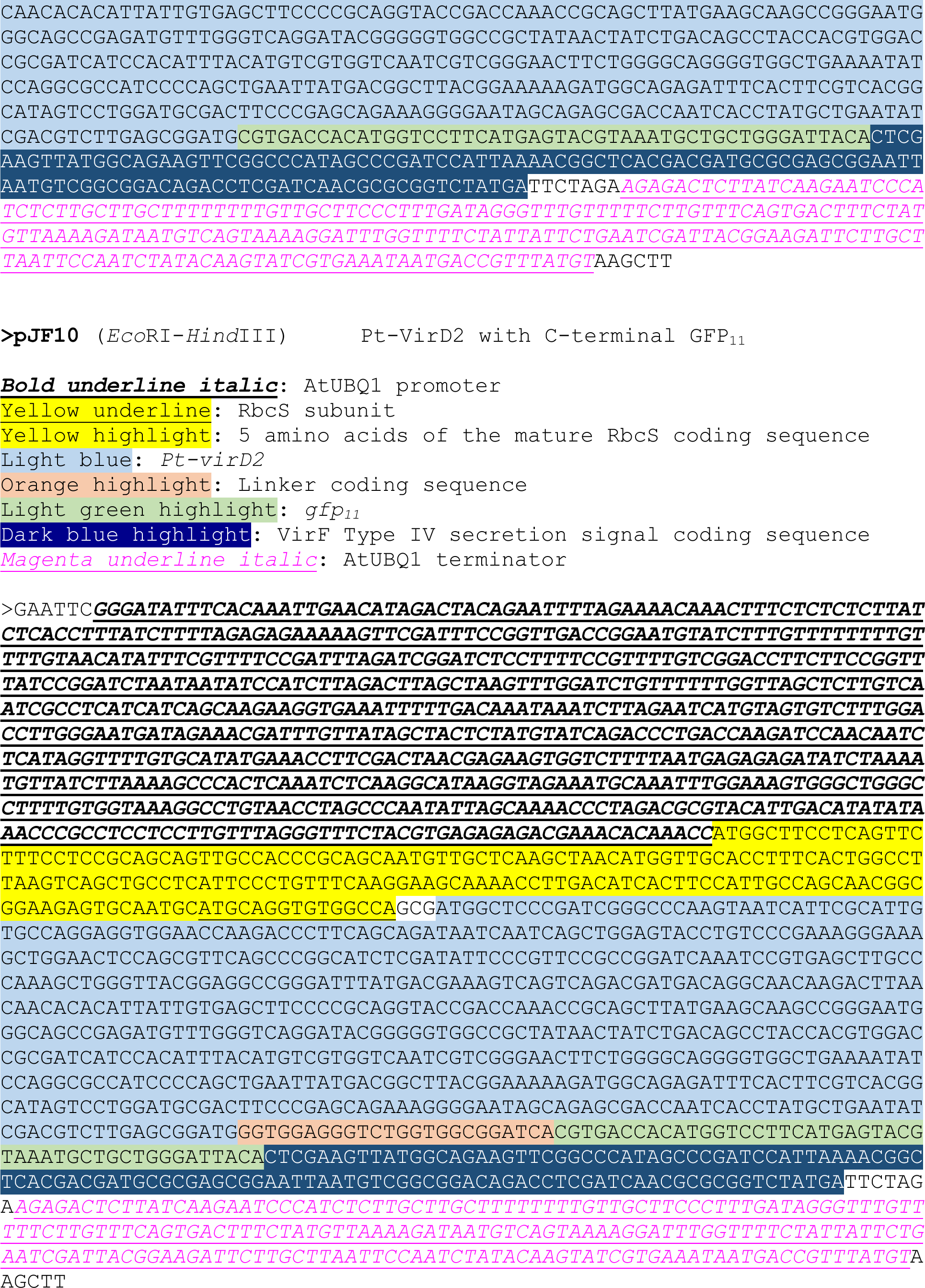
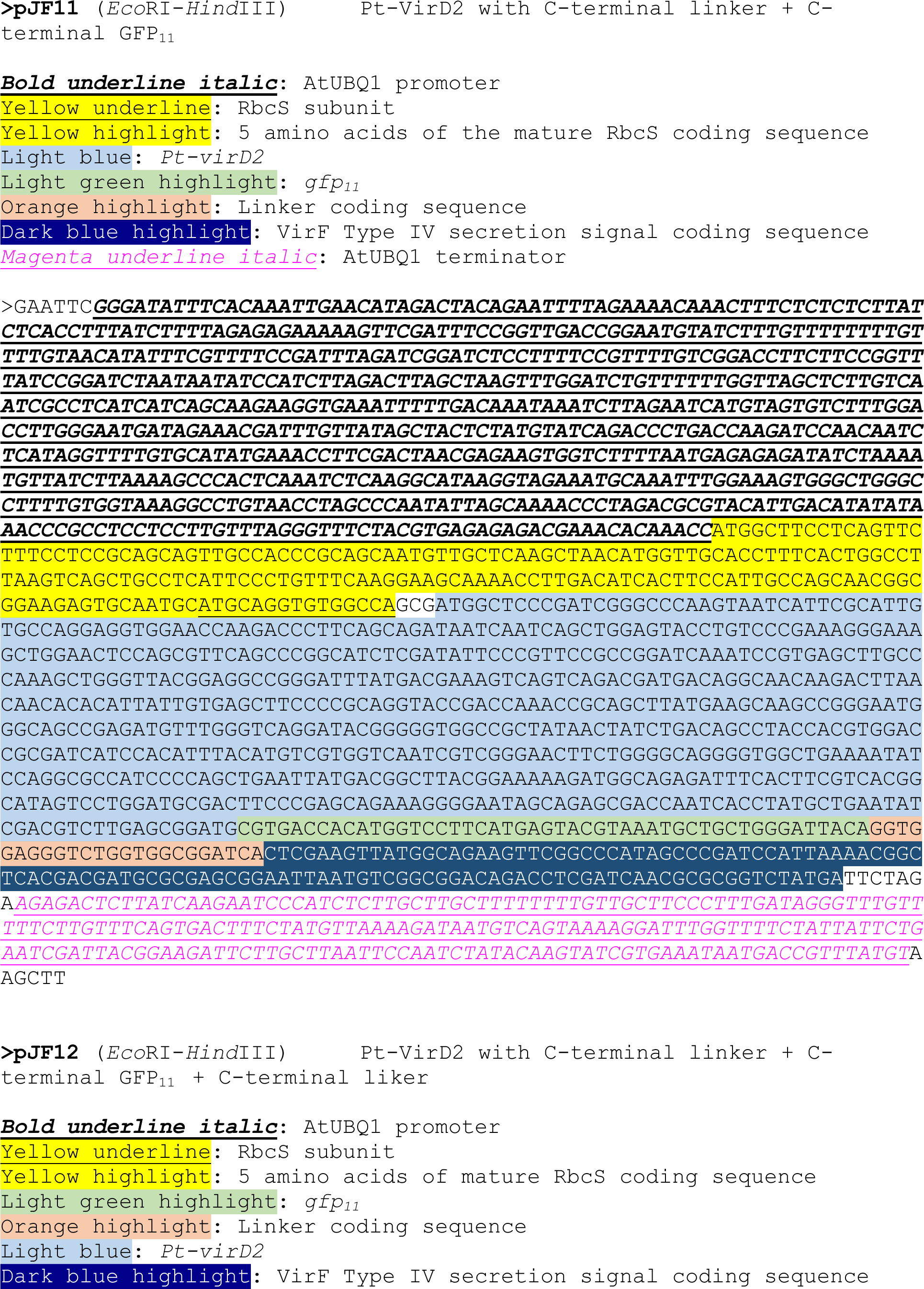
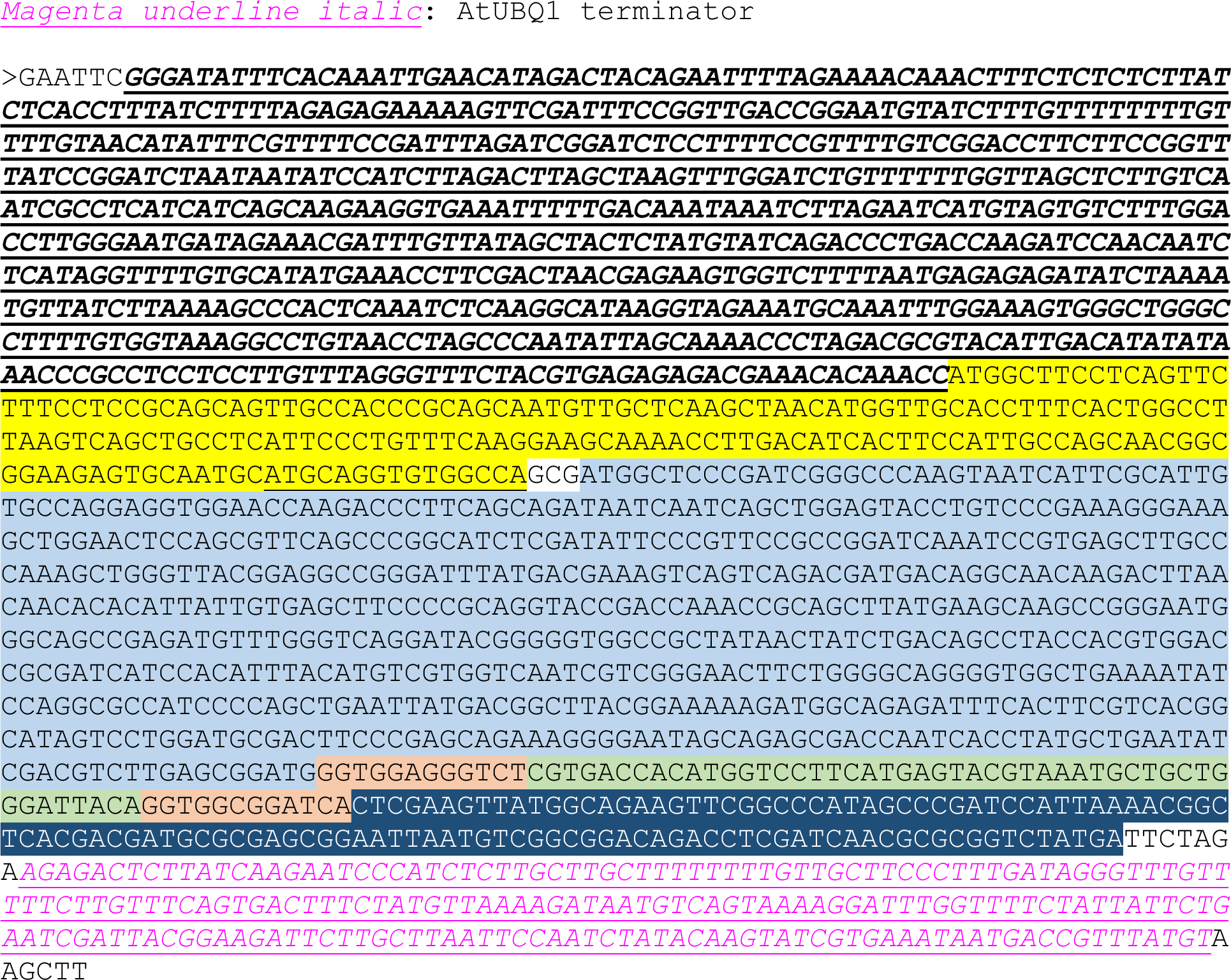
DNA sequence of plasmids used for complementation in the split GFP assay in *E. coli* and in tobacco chloroplasts

